# The E3/E4 ubiquitin ligase UFD-2 mediates negative feedback on Raf protein stability

**DOI:** 10.1101/2022.04.14.488377

**Authors:** Robert Townley, Augustin Deniaud, Kennedy S. Stacy, Claudia S. Rodriguez Torres, Fatemeh Cheraghi, Claire C. de la Cova

## Abstract

Signaling by the kinase cascade comprised of Raf, MEK, and ERK is critical for animal development; moreover, its inappropriate activation is commonly found in human malignancies. In a genetic screen for factors that control signaling by the *Caenorhabditis elegans* Raf ortholog LIN-45, we found that it is negatively regulated by the E3/E4 ubiquitin ligase UFD-2. Both UFD-2 and its partner, the ATP-dependent unfoldase CDC-48, were required for degradation of LIN-45 protein. Our structure-function studies showed that disruption of LIN-45 domains that mediate protein interactions and complex formation, including the Ras binding domain, cysteine-rich domain, or C-terminus, allow for UFD-2-independent degradation. We propose a model whereby UFD-2 mediates a novel step of Raf degradation, by acting with the CDC-48 unfoldase machinery to extract Raf from multiprotein complexes.

**One-Sentence Summary:** Raf kinase complexes are degraded by the UFD-2 ubiquitin ligase and CDC-48 unfoldase during Raf-MEK-ERK signal transduction.

## INTRODUCTION

The Raf family of protein kinases are effectors of Ras signaling that are stimulated by GTP-bound Ras (*1*). Raf directly phosphorylates the kinase MEK, which in turn phosphorylates the MAP kinase ERK. In humans, BRAF, RAF1, and ARAF are Raf family members, while in the nematode *Caenorhabditis elegans*, LIN-45 is the sole ortholog (*2*). Raf proteins share an N-terminal Ras-binding domain (RBD) and cysteine-rich domain (CRD), an intermediate “hinge” region, and a C-terminal kinase domain. When inactive, Raf is a monomer and found in an autoinhibited conformation imposed by interactions between the CRD and 14-3-3 proteins bound to the hinge and C-terminal regions (*3*). Raf activation is stimulated by GTP-Ras binding to the RBD, which relieves it from the autoinhibited conformation and recruits it to the plasma membrane. In this state, Raf proteins dimerize via their kinase domains, allowing for autophosphorylation and activation (*3, 4*). A major mechanism of Raf inactivation involves negative feedback through direct phosphorylation by ERK, resulting hyperphosphorylated, monomeric, and signaling-incompetent Raf (*5*). Among other ERK phosphorylation sites in BRAF is T401 (*6, 7*), a site homologous to T432 in *C. elegans* LIN-45 (*8*). This site is necessary for the function of a Cdc4-phosphodegron (CPD) motif, which is recognized by the E3 ubiquitin ligase SEL-10 or its human ortholog FBXW7. The CPD is required in *C. elegans* LIN-45 and human BRAF for ERK-directed degradation (*8, 9*).

Dysregulated Raf activation is considered a driver event in melanomas, where the activating mutation *BRAF(V600E)* is commonly found in both malignant melanomas and benign, melanocytic nevi (*10*). Despite constitutive kinase activity of BRAF(V600E) protein (*11*), increased ERK activity is relatively rare in nevi carrying *BRAF(V600E)* (*12, 13*), highlighting the importance of additional mutations in melanoma, as well as a need for further knowledge of Raf negative regulation.

In *C. elegans*, Raf signaling impacts multiple developmental and physiological processes (*14*); one example is the patterning of cell fates in the hermaphrodite vulva, involving specification of six epithelial cells termed P3.p-P8.p, or vulval precursor cells (VPCs) (*15*). During the L2 larval stage, production of an EGF-like inductive signal by the Anchor Cell (AC) stimulates EGF receptor (EGFR), Ras, and Raf-MEK-ERK signaling in the nearest VPC, termed P6.p (*16*). In the L3 stage, activation of the ERK ortholog MPK-1 in P6.p promotes a vulval cell fate termed primary (1°). In neighboring cells P5.p and P7.p, activation of Notch and Ral promote an alternative cell fate termed secondary (2°). Signaling by MPK-1 and Notch mutually inhibit one another (*17*); this crosstalk results in a reproducible pattern of 1° and 2° cell fates and ensures the formation of one vulva. Mutations that hyperactivate Ras or Raf signaling result in formation of more than one, a phenotype termed Multivulva (Muv) (*18*).

## RESULTS

To understand the cellular and molecular mechanisms that control signaling by BRAF(V600E), we generated *C. elegans* strains carrying integrated, single-copy transgenes expressing the equivalent mutation, *lin-45(V627E)* specifically in VPCs. A small proportion of adult hermaphrodites carrying *lin-45(V627E)* transgenes displayed the Muv phenotype; however, the majority were wild-type in appearance (Fig. 1A). In a forward genetic screen, we isolated a recessive mutant that displays a highly penetrant Muv phenotype in combination with *lin-45(V627E)*. We mapped and sequenced this mutation and determined this strain carries a nonsense mutation in *ufd-2* (Fig. S1). UFD-2 is the sole *C. elegans* ortholog of human UBE4E, a U-box protein with E3/E4 ubiquitin ligase activities (*19, 20*).

**Fig. 1:**
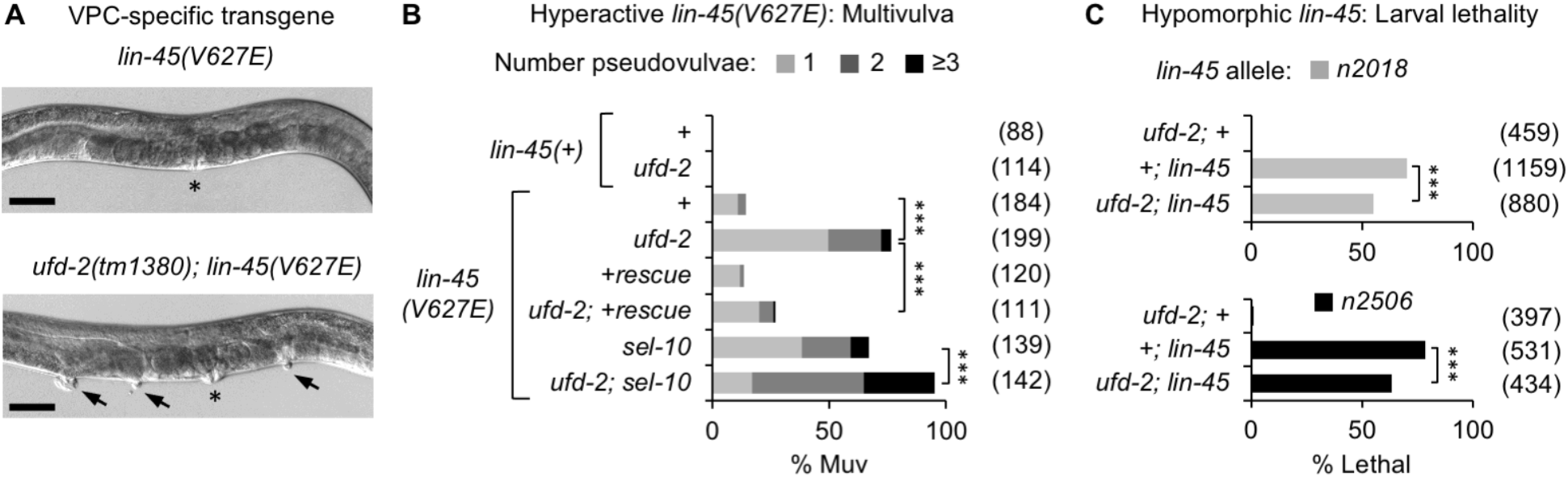
UFD-2 is a negative regulator of LIN-45. **A)** Top; adult hermaphrodite that expresses *lin-45(V627E)*. Bottom; Multivulva (Muv) phenotype of *ufd-2(tm1380); lin-45(V627E)* adult. Asterisk indicates location of vulva; arrowheads indicate pseudovulvae. Scale bar, 50 μm. **B)** Muv phenotype, shown as percentage of total adults. Genotypes were *lin-45(+)* or carried the *covTi106 lin-45(V627E)* transgene. Genotypes rescued for *ufd-2(+)* carried *covTi36* (+rescue). **C)** L1 larval lethality, shown as percentage of total larvae. Genotypes scored were either *ufd-2(0)* or *lin-45* single mutants, or *ufd-2; lin-45* double mutants. The number of animals scored (*n*) is indicated in parentheses. \*\*\**P*<0.0001.

To confirm the role of *ufd-2*, we examined mutants carrying the null allele *ufd-2(tm1380)* (*21*), which we refer to as *ufd-2(0)* throughout this work. For our analysis, we used a *lin-45(V627E)* transgene that caused the Muv phenotype in 14% of adults (Fig. 1B). *ufd-2(0)* mutants were not Muv in the presence of *lin-45(+)*; however, *ufd-2(0)* mutants carrying the *lin-45(V627E)* transgene displayed a highly penetrant Muv phenotype affecting 76% of adults (Fig. 1A,B). The increased penetrance was reverted by the introduction of a single-copy *ufd-2(+)* transgene expressed specifically in VPCs (Fig. 1B), confirming that *ufd-2* is responsible for the Muv phenotype and that it acts cell-autonomously in VPCs. As previously reported, the Muv phenotype of *lin-45(V627E)* animals was enhanced by the *sel-10(ok1632)* null mutation, referred to here as *sel-10(0)* (Fig. 1B) (*8*). In animals lacking both *ufd-2* and *sel-10*, penetrance of the Muv phenotype in *lin-45(V627E)* animals was significantly increased compared to either single mutant (Fig. 1B), suggesting that UFD-2 and SEL-10 function in parallel for some roles.

If UFD-2 inhibits LIN-45 activity, then loss of *ufd-2* may suppress phenotypes resulting from reduced LIN-45 activity. We examined two alleles of *lin-45* that alter the LIN-45 RBD and result in partial loss of function and lethality at the L1 larval stage (*22, 23*). Compared to either *lin-45* mutant alone, the *ufd-2(0); lin-45* double mutants displayed a significant reduction in lethality (Fig. 1C). These results support a role for UFD-2 in negatively regulating LIN-45 and exclude the possibility that the regulation is limited to vulval induction.

### Negative regulation of Raf-MEK-ERK activity

In VPCs, patterned fate specification is initiated by an inductive signal from the AC (*18*). Consequently, activated MPK-1 phosphorylates the Elk1 ortholog LIN-1 (*24*), resulting in derepression of LIN-1 targets in P6.p, including the Delta-like gene *lag-2* (*25*) (Fig. 2A). The transgene *arIs222* contains a *lag-2* promoter fragment and drives expression of 2xNLS-tagRFP exclusively in VPCs that adopt 1° cell fate, i.e. P6.p and its descendants as VPCs undergo cell divisions (*26*). In genotypes with increased Raf signaling, this reporter is expressed ectopically in additional VPCs (*27*). To ensure that fate specification events were completed, we scored VPCs that had completed one or two cell divisions, referring to the descendants with a suffix; for example, descendants of P6.p were termed “P6.pxx.” Loss of *ufd-2* alone did not alter the P6.pxx expression pattern of the *lag-2* reporter (Fig. 2B). Expression of activated *lin-45(V627E)* caused ectopic reporter expression in a significant portion of animals, typically in descendants of P4.p and P8.p, a pattern consistent with previous observations (*28*). In those expressing *lin-45(V627E)*, loss of *ufd-2* enhanced the frequency of ectopic *lag-2* reporter expression (Fig. 2B), suggesting that UFD-2 acts to reduce MPK-1 activity and subsequent expression of *lag-2* in VPCs.

**Fig. 2:**
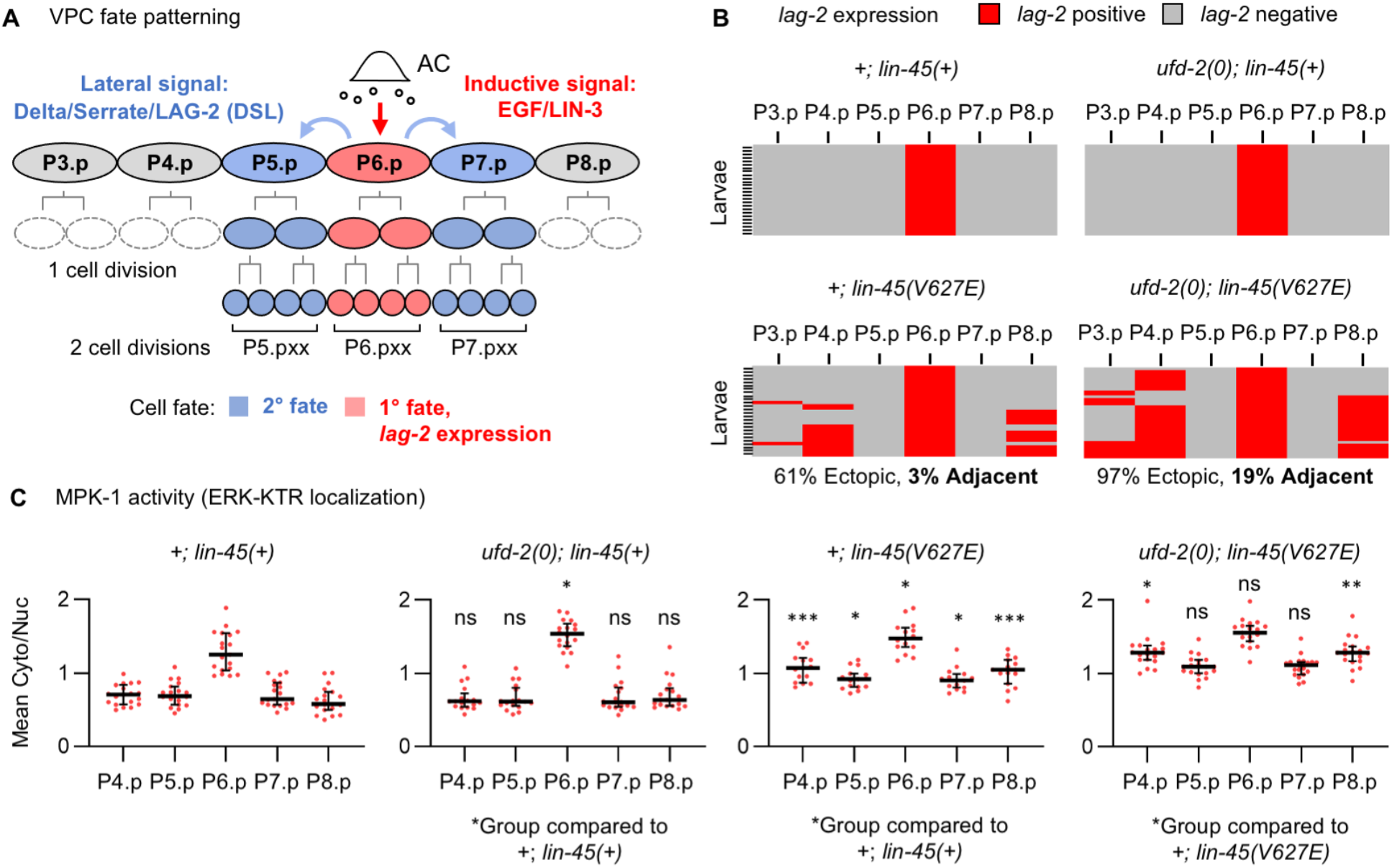
UFD-2 negatively regulates LIN-45 signaling. **A)** An inductive signal activates Raf-MEK-ERK signaling in P6.p, promoting *lag-2* expression and 1° fate (red). A lateral signal promotes 2° fate (blue) in P5.p and P7.p. **B)** Expression of a *lag-2* reporter, shown as *lag-2*-positive (red) or *lag-2*-negative (gray). In all panels, rows represent individual L3 stage larvae; columns represent descendants of VPCs. Genotypes were *+; lin-45(+)* (n=37), *ufd-2(0); lin-45(+)* (n=36), *+; lin-45(V627E)* (n=31), or *ufd-2(0); lin-45(V627E)* (n=36). **C)** MPK-1 activation in individual VPCs assessed using ERK-KTR localization. Data points represent the mean Cyto/Nuc ratio for individual cells. Genotypes scored were *+; lin-45(+)* (n=19), *ufd-2(0); lin-45(+)* (n=18), *+; lin-45(V627E)* (n=15), or *ufd-2(0); lin-45(V627E)* (n=19). Reported *P* values were adjusted for multiple comparisons; ns not significant, \**P* <0.05, \*\**P* <0.001, \*\*\**P* <0.0001.

To more directly test how loss of *ufd-2* impacts MPK-1 activity, we made use of ERK-KTR, a Kinase Translocation Reporter (KTR) that responds to phosphorylation by MPK-1 through a change in its nuclear-cytoplasmic localization (*16, 29*). In the presence of active MPK-1, ERK-KTR is localized to the cytoplasm; in the absence of MPK-1 activity, ERK-KTR accumulates in the nucleus. We examined ERK-KTR during the L3 stage before VPC divisions, a time when MPK-1 activation is robust in P6.p (*16*). In wild-type animals, MPK-1 activation was greatest in P6.p (Fig. 2C). This pattern was maintained in *ufd-2(0)* mutants; however, MPK-1 activation in P6.p was significantly increased. In animals expressing *lin-45(V627E)*, MPK-1 activation was elevated in all VPCs. Compared to the *lin-45(V627E*) strain, MPK-1 activation in *ufd-2(0)*; *lin-45(V627E)* double mutants was further increased in P4.p and P8.p (Fig. 2C), a pattern similar to the ectopic *lag-2* reporter expression (Fig. 2B). These results show that loss of *ufd-2* function results in greater MPK-1 activation and that UFD-2 inhibits signaling by both the LIN-45(+) and LIN-45(V627E) forms.

### LIN-45 protein degradation

To test whether *ufd-2* loss impacts LIN-45 protein degradation, we generated N- and C-terminal GFP knock-in alleles (Fig. S2A,B) and examined the expression of tagged protein in P5.pxx, P6.pxx, and P7.pxx cells. When tagged at the either the N-terminus (Fig. 3A) or C-terminus (Fig. S2D), LIN-45 protein is visible in P5.pxx and P7.pxx, and noticeably lower in P6.pxx. Loss of *ufd-2* increased the visible protein accumulation of GFP-LIN-45 in P6.pxx (Fig. 3A). Increased accumulation was seen in 68% of *ufd-2(0)* mutants, a significantly higher portion than in wild type (Fig. 3B). Loss of *sel-10* increased the accumulation of GFP-LIN-45 seen in P6.pxx (Fig. S2C) in 100% of *sel-10(0)* mutants (Fig. 3B).

**Fig. 3.**
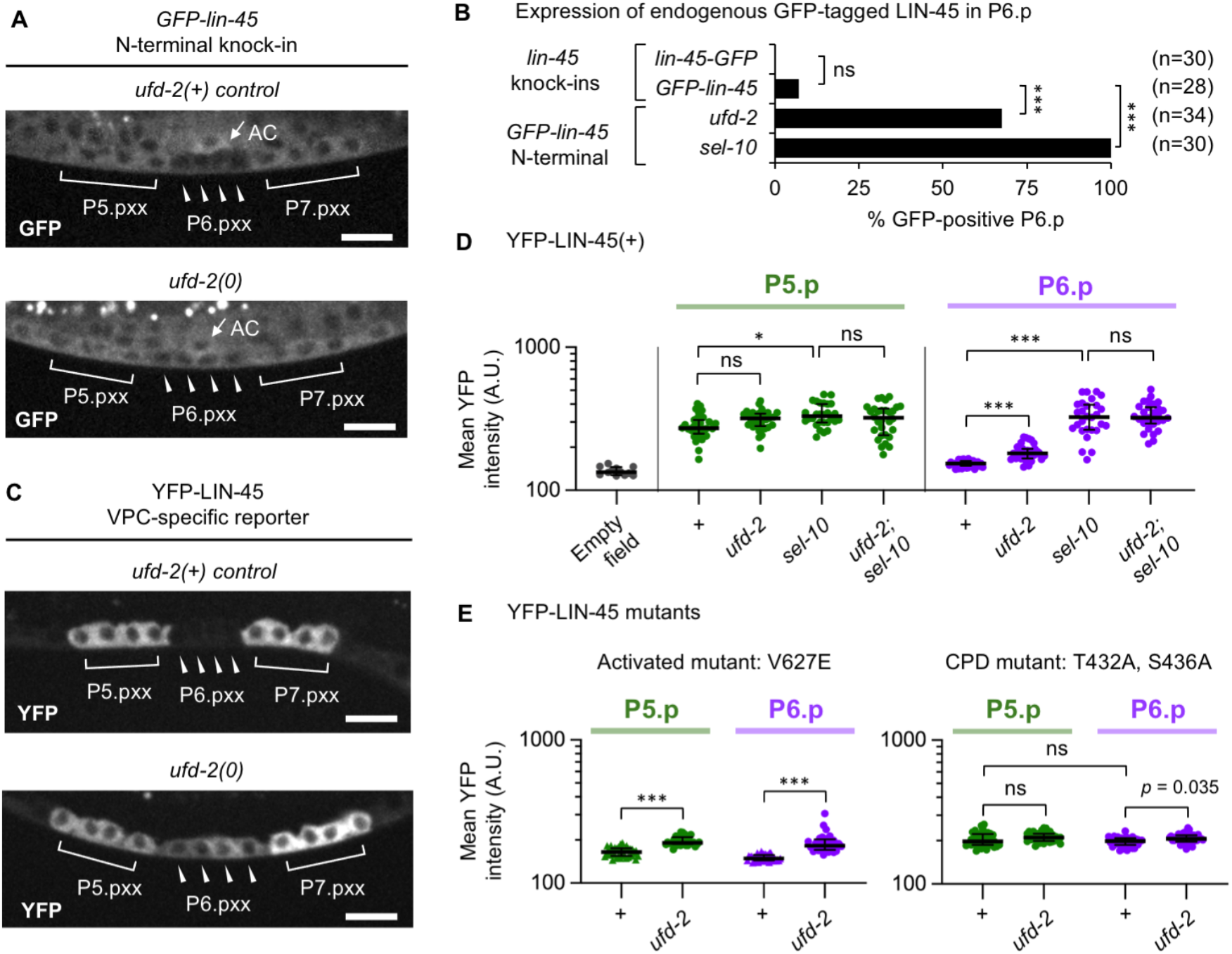
UFD-2 is required for LIN-45 protein degradation. **A)** Expression of the *GFP-lin-45* knock-in. Shown are L3 larval stage P5.pxx, P6.pxx, and P7.pxx cells in wild type and *ufd-2(0)* mutants; the anchor cell (AC). Scale bar, 10 μm. **B)** Expression of GFP-LIN-45 in P6.pxx cells, shown as percentage of animals in wild type, *ufd-2(0)*, or *sel-10(0)* genotypes. **C)** Expression of the VPC-specific reporter YFP-LIN-45 in wild type and *ufd-2(0)* mutants. **D-E)** Intensity of YFP-LIN-45 in P5.pxx (green) and P6.pxx (purple). Mean YFP intensities were plotted on a log_10_ scale. D) YFP-LIN-45(+) intensity in wild type (n=32), *ufd-2(0)* (n=33), *sel-10(0)* (n=28), or *ufd-2(0); sel-10(0)* (n=30). E) At left, YFP-LIN-45(T432A, S436A) intensity in wild type (n=35) or *ufd-2(0)* (n=37). At right, YFP-LIN-45(V627E) intensity in wild type (n=36) or *ufd-2(0)* (n=32). Reported *P* values were adjusted for multiple comparisons; ns not significant, \**P* <0.01, \*\*\**P* <0.0001.

To quantify LIN-45 protein levels, we made use of an integrated, single-copy *yfp-lin-45* transgene that expresses specifically in VPCs. Our transgene employs regulatory elements from the *lin-31* gene (referred to here as *lin-31p*); when used to drive expression of YFP alone, it is observed in all six VPCs and their descendants throughout the L3 stage. When used to express YFP-tagged *lin-45*, YFP-LIN-45 protein is downregulated post-transcriptionally in P6.p at the L3 stage, (*8*) (Fig. 3C). In addition to recapitulating the protein pattern seen in our knock-in alleles, the *lin-31p::yfp-lin-45* transgene fully rescued vulval development of a *lin-45* null mutant (Fig. S2E). All additional YFP-LIN-45 reporters were made using the same strategy. We measured YFP fluorescence intensity in two contexts: P5.pxx, where MPK-1 signaling is inactive, and P6.pxx, where it is active. In *ufd-2(0)* mutants, the YFP-LIN-45 level in P5.pxx was unchanged compared to wild type; however, YFP-LIN-45 in P6.pxx was visibly higher and significantly increased compared to wild type (Fig. 3C,D). We next tested whether *ufd-2* impacts the degradation of YFP-LIN-45(V627E). Loss of *ufd-2* significantly elevated the intensity of YFP-LIN-45(V627E) in both P5.pxx and P6.pxx (Fig. 3E).

Because SEL-10 acts in negative feedback on LIN-45, we tested whether loss of *ufd-2* and *sel-10* are additive with respect to LIN-45 protein. As previously shown, YFP-LIN-45 intensity was significantly increased in P6.pxx of *sel-10(0)* mutants (Fig. 3D) (*8*). However, YFP-LIN-45 levels were no greater in the *ufd-2(0); sel-10(0)* double mutants. LIN-45 CPD phosphorylation is also required for negative feedback by ERK and SEL-10, and is disrupted by mutation of residues T432 and S436 (*8*). In *ufd-2(0)* mutants, we found no additional increase in the intensity of the reporter YFP-LIN-45(T432A, S436A) bearing these mutations (Fig. 3E).

In *C. elegans*, UFD-2 can cooperate with chaperones and the chaperone-associated E3 ubiquitin ligase CHN-1, an ortholog of human STUB1, to promote polyubiquitination (*19, 21*). Additionally, UFD-2 interacts with CDC-48 (*30*), ortholog of the Cdc48/p97 protein unfoldase. To understand the mechanistic details of UFD-2-mediated regulation, we assessed the effect of *chn-1* mutation and *cdc-48* knock-down on LIN-45 degradation.

The viable allele *chn-1*(*by155)* is a deletion that eliminates the C-terminal U-box required for E3 ligase activity and binding to UFD-2 (*21*). We assessed YFP-LIN-45 degradation in this mutant and found there was no effect on YFP-LIN-45 intensity in P6.pxx (Fig. S3A).

We used RNAi to deplete *cdc-48*.*1* and *cdc-48*.*2*, two highly similar genes encoding *C. elegans* orthologs (*31*). To avoid the embryonic lethality of *cdc-48*.*1* and *cdc-48*.*2* depletion (*32*), animals were treated after embryogenesis and examined at the L3 larval stage. Compared to animals treated with *lacZ* RNAi as a negative control, expression of YFP-LIN-45 was significantly increased in P6.pxx of those depleted for both *cdc-48*.*1* and *cdc-48*.*2* simultaneously (Fig. 4A). *cdc-48*.*1/2* RNAi-treated animals failed to develop later; however, VPC fate specification at the L3 stage appeared to be normal, as P5.p, P6.p, and P7.p all completed two cell divisions.

**Fig. 4.**
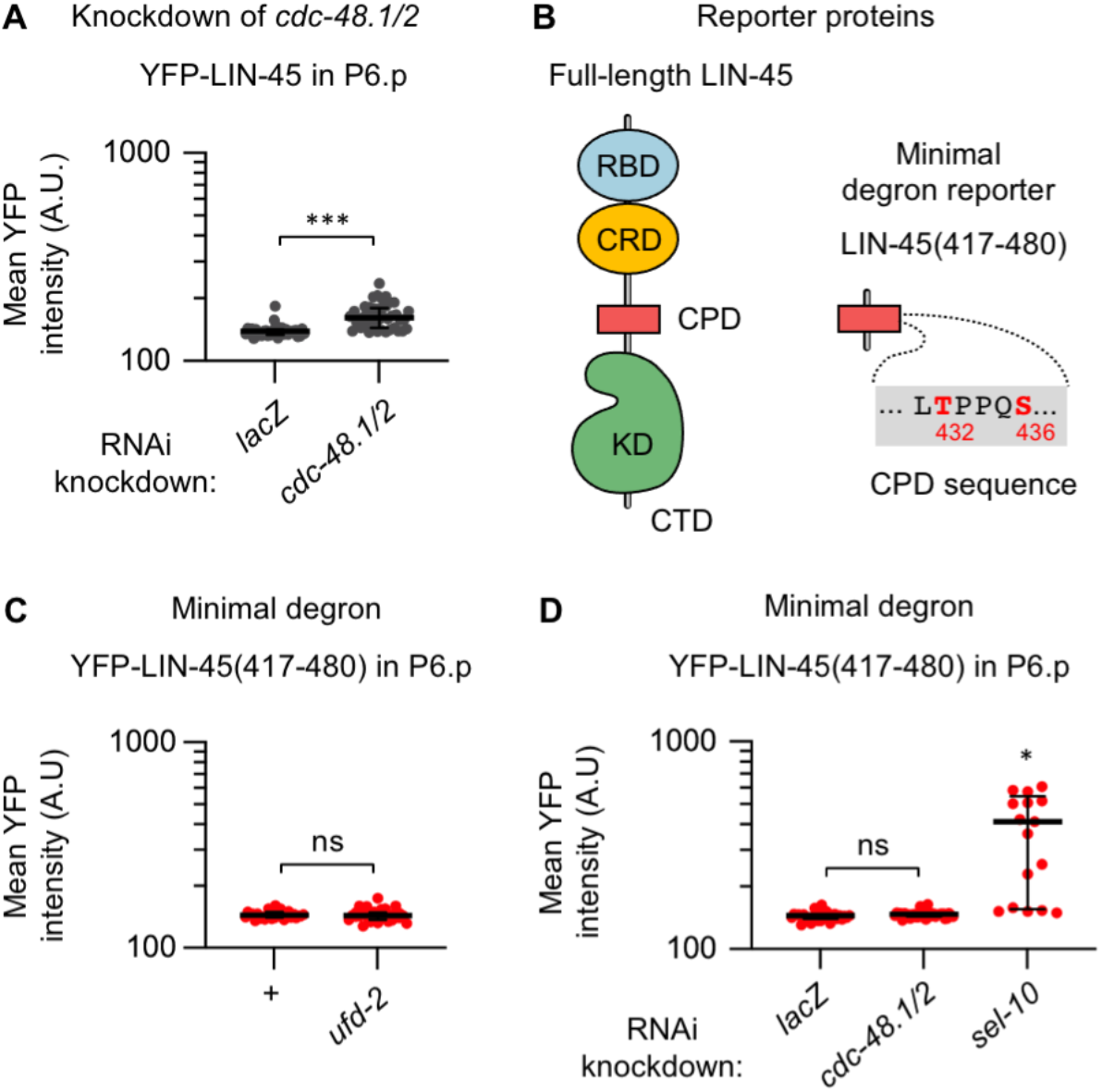
LIN-45 degradation requires the CDC-48 unfoldase. **A)** Expression of YFP-LIN-45(+) in P6.pxx cells of animals treated with RNAi targeting a control (*lacZ*) (n=33) or the genes *cdc-48*.*1* and *cdc-48*.*2* (n=42). For all panels, mean YFP intensities are plotted on a log_10_ scale. **B)** Domains of LIN-45 include the RBD (blue), CRD (yellow), CPD (red), kinase domain (KD, green), and C-terminal tail (CTD). The minimal degron of LIN-45 contains residues 417-480 and a canonical CPD. **C)** Expression of YFP-LIN-45(417-480) in P6.pxx of wild type (n=31) and *ufd-2(0)* (n=31). **D)** Expression of YFP-LIN-45(417-480) in P6.pxx of animals treated with *lacZ* RNAi (n=31), or *cdc-48*.*1/2* RNAi (n=36), or *sel-10* RNAi (n=17). Reported *P* values were adjusted for multiple comparisons; ns not significant, \**P* <0.01, \*\*\**P* <0.0001.

Substrates of Cdc48 are often well-folded, or part of multiprotein complexes, or localized to membranes. By unfolding such proteins, Cdc48 is thought to prepare them for degradation by the proteasome (*33*). If LIN-45 protein requires this processing, we reasoned that an unfolded substrate derived from LIN-45 would bypass the requirements for *ufd-2* and *cdc-48*. To test this prediction, we used a reporter expressing a YFP-tagged region of LIN-45 termed the “minimal degron,” or YFP-LIN-45(417-480) (Fig. 4B) (*28*). This short substrate contains the CPD motif and is degraded in P6.pxx (Fig. S3B). In contrast to full-length protein, the minimal degron reporter was not stabilized in *ufd-2(0)* mutants (Fig. 4C), or in animals depleted of *cdc-48*.*1* and *cdc-48*.*2* (Fig. 4D). However, this reporter was stabilized by depletion of *sel-10* (Fig. 4D), confirming its degradation still relies on SEL-10. These results suggest that native, full-length LIN-45 contains structural features that impede degradation in the absence of UFD-2 and CDC-48 activities.

### LIN-45 domains required for UFD-2 dependence

To identify the structural features that make LIN-45 degradation dependent on UFD-2, we systematically perturbed LIN-45 domains and protein-protein interactions by mutation. LIN-45 contains several domains present in all three human Raf proteins (Fig. 5A). Among these are the RBD, the CRD, a 14-3-3 binding site located within the “hinge” region, the C-terminal kinase domain, and a second 14-3-3 binding site located near the C-terminus (*1*).

**Figure 5.**
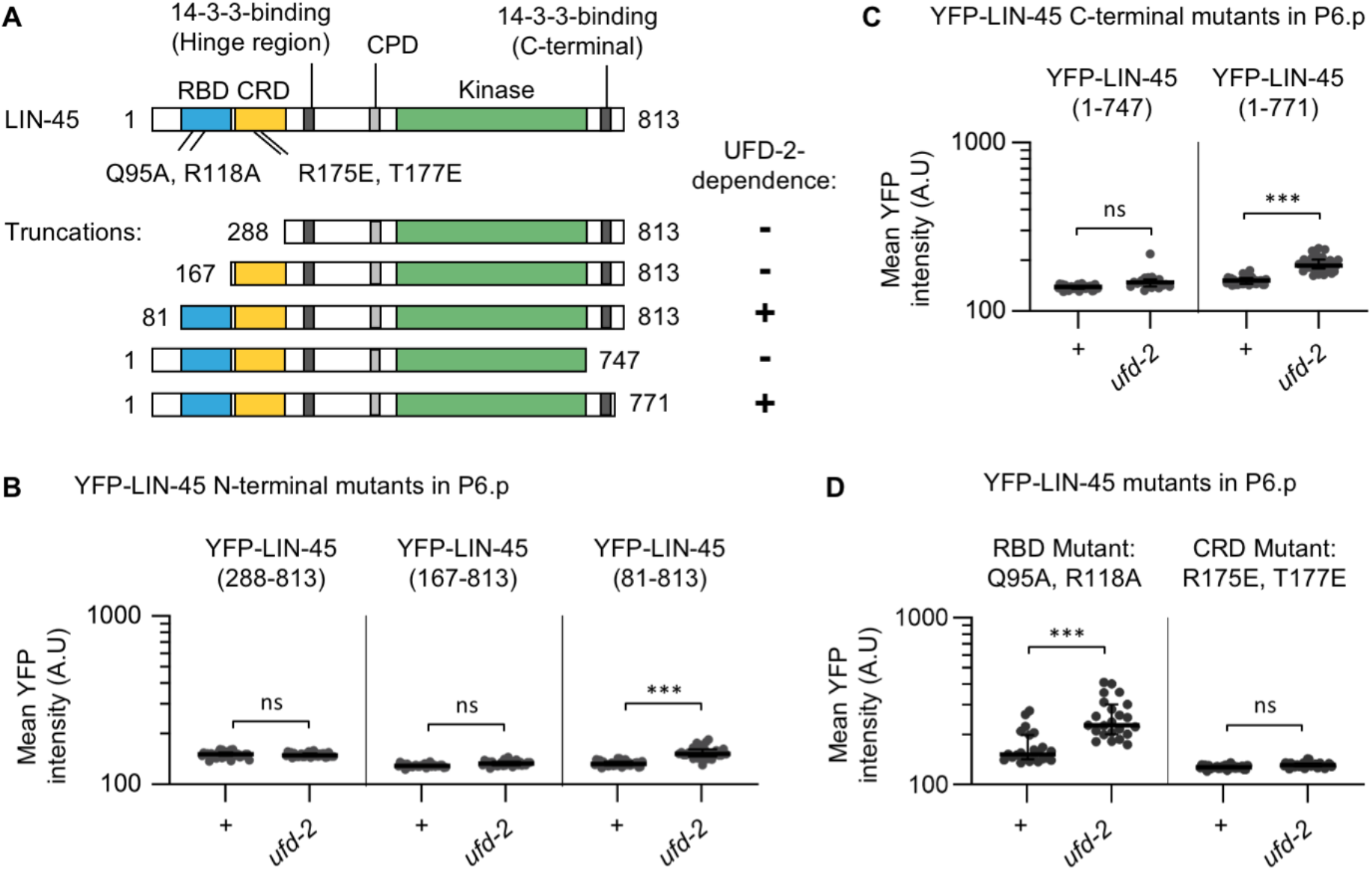
LIN-45 mutants bypass the requirement for UFD-2. Mutant forms of YFP-LIN-45 were examined in wild type and *ufd-2(0)* mutants. **A)** LIN-45 protein and domains. Shown are locations of mutations and truncations. At right, whether protein degradation in P6.pxx cells was UFD-2-dependent (+) or independent (-) is indicated. **B-D)** Protein levels of YFP-LIN-45 mutant reporters in P6.pxx. For all panels, mean YFP intensities are plotted on a log_10_ scale. B) N-terminal truncations were: YFP-LIN-45(288-813), in wild type (n=36) and *ufd-2(0)* (n=32); YFP-LIN-45(167-813), in wild type (n=28) and *ufd-2(0)* (n=29); YFP-LIN-45(81-813), in wild type (n=31) and *ufd-2(0)* (n=36). C) C-terminal truncations were: YFP-LIN-45(1-747), in wild type (n=31) and *ufd-2(0)* (n=19); YFP-LIN-45(1-771), in wild type (n=35) and *ufd-2(0)* (n=36). **C)** Missense mutations scored were YFP-LIN-45(Q95A, R118A), in wild type (n=23) and *ufd-2(0)* (n=23), and LIN-45(R175E, T177E) in wild type (n=29) and *ufd-2(0)* (n=40). Reported *P* values were adjusted for multiple comparisons; ns not significant, \**P* <0.01, \*\*\**P* <0.0001.

We tested the roles of LIN-45 N-terminal domains by expressing forms truncated at three points within the N-terminus (Fig. 5A). Two of these forms, YFP-LIN-45(288-813) and YFP-LIN-45(167-813), disrupt or remove the RBD and CRD. Both were degraded in P6.pxx in wild type and *ufd-2(0)* mutants (Fig. 5B), indicating that they bypass the requirement for UFD-2. One longer form, YFP-LIN-45(82-813), retains both the RBD and CRD. This form was degraded in P6.pxx in wild type and stabilized in *ufd-2(0)* mutants (Fig. 5B), indicating that it retains UFD-2 dependence. We tested the role of the LIN-45 C-terminus by expressing two truncations (Fig. 5A). YFP-LIN-45(1-747) lacks all sequences C-terminal to the kinase domain. This form was degraded in P6.pxx in wild type and *ufd-2(0)* mutants (Fig. 5C), indicating that it bypasses UFD-2. In contrast, YFP-LIN-45(1-771) retains the C-terminal 14-3-3 binding site; this form was degraded in wild type and stabilized in *ufd-2(0)* mutants (Fig. 5C), indicating that it is dependent on UFD-2.

To further define the roles of N-terminal domains, we generated missense mutations in full-length YFP-LIN-45. One possibility is that Ras-binding confers UFD-2 dependence, possibly by recruiting LIN-45 to the plasma membrane. We tested this by creating mutations at Q95 and R118 (Fig. S4), residues required for Ras-binding in vitro and LIN-45 function in vivo (*23, 34*). The mutant YFP-LIN-45(Q95A, R118A) was degraded in P6.pxx in wild type and stabilized in *ufd-2(0)* mutants (Fig. 5D), suggesting that Ras-binding is not important for dependence on UFD-2. It should be noted that this transgene is expressed in animals with a wild-type *lin-45(+)* allele and vulval induction is normal. We found that the YFP-LIN-45(Q95A, R118A) form was incapable of rescuing vulval development in the *lin-45* null mutant, confirming that the RBD mutations indeed abrogate function (Fig. S2E).

A second N-terminal domain implicated in UFD-2 dependence is the CRD. To test its function, we generated mutations in R175 and T177 (Fig. S4), conserved residues that make contact with 14-3-3 proteins and Ras (*35, 36*). The mutant YFP-LIN-45(R175E, T177E) was degraded in P6.pxx in both wild type and *ufd-2(0)* mutants (Fig. 5D), implicating protein interactions mediated by this domain in UFD-2 dependence. Because the CRD interacts with i) 14-3-3 proteins in the autoinhibited form and ii) Ras dimers and phospholipids in the active form, its mutation may disrupt LIN-45 complex formation or recruitment to the plasma membrane. We found that the YFP-LIN-45(R175E, T177E) mutant reporter partially rescued normal vulval induction in the *lin-45* null (50% of animals, Fig. S2E). Strikingly, another 25% were Vulvaless, indicating no induction, while 25% had a Muv phenotype characteristic of ectopic induction (Fig. S2E), results consistent with roles for the CRD in both autoinhibition and activation.

Our data from mutant YFP-LIN-45 reporters support the possibility that efficient LIN-45 degradation relies on UFD-2 because of LIN-45 protein-protein interactions. Specifically, we find evidence that interactions at the RBD and CRD, but not Ras-binding *per se*, and interactions at the C-terminal 14-3-3 protein binding site, are required for UFD-2 dependence.

## DISCUSSION

We identified the E3/E4 ubiquitin ligase UFD-2 and the unfoldase CDC-48 as negative regulators of LIN-45 protein stability and Raf-MEK-ERK signaling. In this discussion, we address how UFD-2 function relates to ERK-directed negative feedback, briefly summarize known UFD-2 and CDC-48 molecular activities, and propose a model whereby these factors are needed to unfold and extract LIN-45 from multiprotein complexes (Fig. 6).

**Fig. 6.**
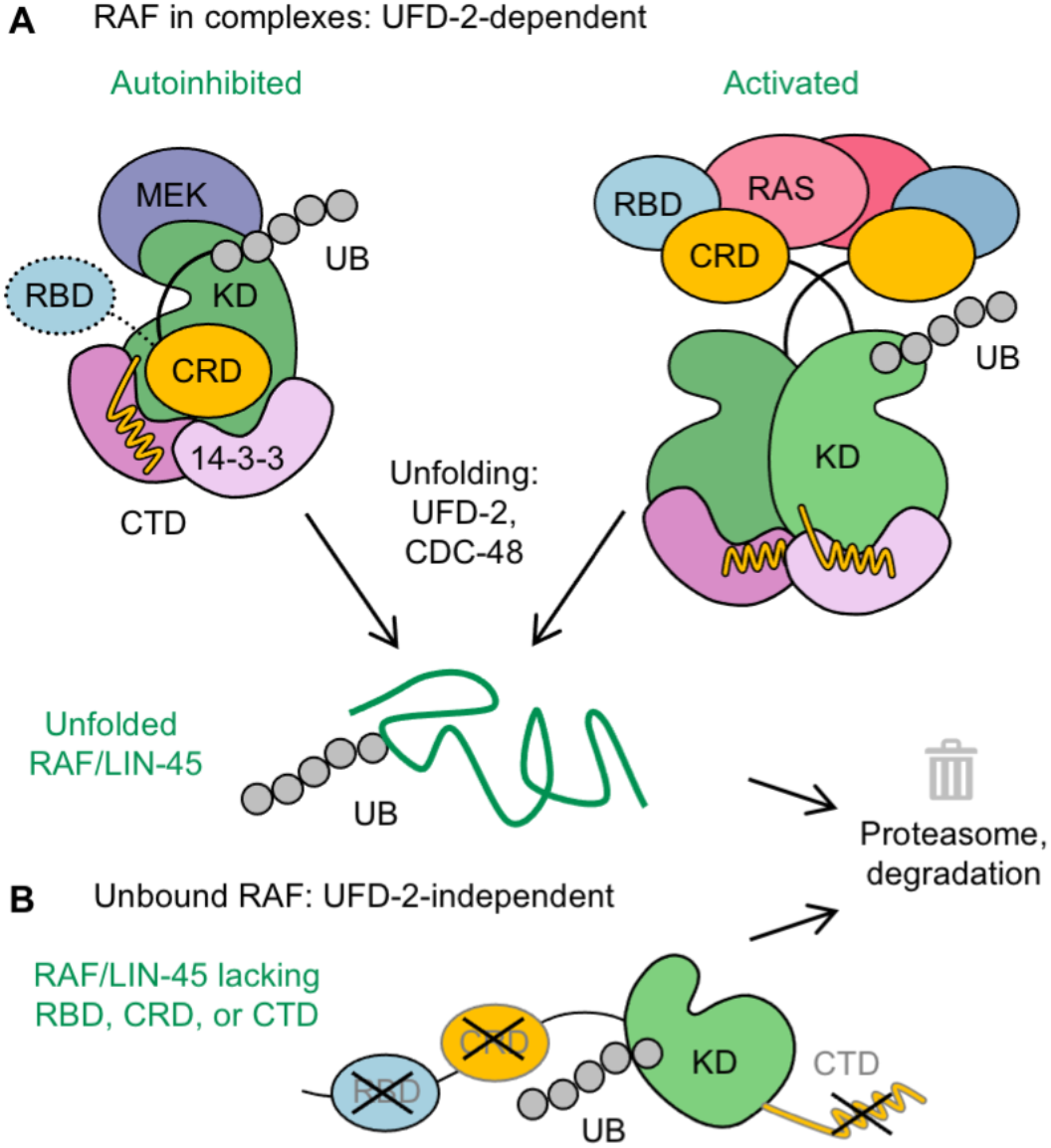
UFD-2 is required for degradation of LIN-45 in multiprotein complexes. **A)** The known Raf/LIN-45 protein complexes mediated by the Ras-binding domain (RBD, blue), cysteine-rich domain (CRD, yellow), kinase domains (KD, green), and C-terminal domain (CTD, yellow line). In the autoinhibited state, Raf/LIN-45 binds 14-3-3 proteins via its CRD, CTD, and hinge region. In the activated dimer, Raf/LIN-45 binds RAS via its RBD and CRD, and interacts with 14-3-3 proteins via the CTD. We propose that proteasome targeting of LIN-45 is impeded by multiprotein complexes; when unfolded via the activities of UFD-2 and CDC-48, LIN-45 is degraded. **B)**. LIN-45 mutants lacking RBD, CRD, or CTD functions are degraded independently of UFD-2.

### Negative feedback and LIN-45 degradation

In human cells, ERK activation stimulates Raf phosphorylation (*5, 7*) and in some contexts, its degradation (*9*). In *C. elegans*, the spatial and temporal pattern of LIN-45 degradation in VPCs is controlled by the kinases MPK-1, GSK-3 and CDK-2, and the E3 ubiquitin ligase SEL-10 (*28*). We propose that UFD-2 promotes LIN-45 degradation in the same negative feedback pathway as SEL-10. With respect to LIN-45 protein levels, two genetic epistasis results support this model. First, LIN-45 levels in a *ufd-2(0)*; *sel-10(0)* double mutant were unchanged from those in a *sel-10* mutant. Secondly, loss of *ufd-2* failed to further stabilize LIN-45 protein that lacks a functional CPD. It is possible that both ubiquitin ligases act directly on LIN-45; indeed, both are capable of efficiently generating polyubiquitin chains (*19, 37*). A major difference between the two lies in their substrate targeting mechanisms; while SEL-10 directly binds a CPD motif within its substrates, UFD-2 is recruited through association with chaperones or CDC-48, as described later in this discussion. In contrast to the non-additive effect on LIN-45 protein, we observed an additive effect of *ufd-2(0)*; *sel-10(0)* double mutants on the Muv phenotype, suggesting that UFD-2 or SEL-10 may have additional non-overlapping substrates that impact cell fate specification in VPCs.

A consequence of UFD-2 acting in ERK-directed negative feedback is that its loss specifically impacts cells where MPK-1 activity is high. We observed this in wild-type VPCs, where loss of *ufd-2* increased LIN-45 protein level only in P6.pxx cells, and in VPCs expressing *lin-45(V627E)*, where loss of *ufd-2* also affected P5.pxx. Quantitation of LIN-45 protein levels in P6.pxx cells allowed us to estimate that 25% of LIN-45 protein remained stable in the absence of UFD-2 (compared to the protein levels in P6.pxx in *sel-10(0)*) (Fig. 3D). It is possible that UFD-2 is required for degradation of a subset of LIN-45 proteins, perhaps due to their physical structure, protein interactions, localization, or post-translational modifications.

### Roles of UFD-2 and CDC-48

Studies of yeast and human Cdc48/p97 protein show that it acts as an ATP-dependent unfoldase that facilitates degradation of polyubiquitinated proteins (*38*). Cryo-electron microscopy reveals that Cdc48 forms a hexamer ring of ATPase domains surrounding a central pore (*33, 39*). A ubiquitinated substrate is recruited by adaptors Ufd1 and Npl4, and the polypeptide unfolded during translocation through the pore. A required step for release is trimming of a substrate’s polyubiquitin chains, leaving many with fewer than ten ubiquitin moieties (*40*). However, the Cdc48 complex also interacts with Ufd2, which acts to extend the ubiquitin chains of Cdc48 substrates (*41*). One model of Cdc48-Ufd2 function posits that Ufd2 is responsible for polyubiquitination of newly trimmed substrates as they exit the Cdc48 complex, ensuring their efficient transfer to the proteasome (*38*).

We propose that UFD-2 and CDC-48 are required for the degradation of ubiquitinated LIN-45 proteins that participate in multiprotein complexes, e.g. in the autoinhibited monomer and/or activated dimers (Fig. 6A). Furthermore, “unbound” LIN-45 may be available to the proteasome without processing by UFD-2 and CDC-48; our data on mutant forms of LIN-45 are informative about which protein-protein interactions cause UFD-2 dependence (Fig. 6B).

Mutations at the C-terminus of YFP-LIN-45 implicate the region between residues 747 and 771 in UFD-2 dependence. This region is highly conserved in human Raf proteins (Fig. S4B); it contains a phosphorylation-regulated motif that interacts with 14-3-3 (*3, 4*), and is required for maximal kinase activity (*42*). We propose that LIN-45 protein bound to 14-3-3 at this site requires the activities of UFD-2 and CDC-48 for efficient degradation. It should be noted that an unstructured C-terminus may allow some substrates to bypass Cdc48 (*43*).

However, we think this is unlikely to account for independence of the shorter truncation LIN-45(1-747), because it terminates at a well-ordered endpoint identical to BRAF kinase domains expressed in crystal structures (*44*).

Mutations at the N-terminus of YFP-LIN-45 implicate the RBD and CRD regions in UFD-2 dependence. For example, mutation of conserved residues R175 and T177 in the CRD (Fig. S4A) allowed degradation independently of UFD-2. Structural studies suggest these residues impact incorporation in the activated dimer, or in a second possibility that is not mutually exclusive, interactions with 14-3-3 in the autoinhibited state (*3, 36*). Mutations in the RBD were also informative; surprisingly, its deletion allowed UFD-2 independence, while missense mutations did not. We infer that RBD mediates LIN-45 interactions with factors other than Ras. One candidate function is assisting incorporation into the very large “signalosome” at the plasma membrane, where Raf interacts with Ras and Galectin proteins (*36, 45*). Finally, after activation, Raf also transitions to a high molecular weight complex where it is bound by the chaperones HSP90 and CDC37 (*46*). Due to their large size, complexity, or localization, it is possible that these structures may impede proteasome targeting and degradation.

## MATERIALS AND METHODS

### C. elegans *genetics*

The complete genotypes of *C. elegans* strains used in this work are listed in Table S1. The following alleles were obtained from the Caenorhabditis Genetics Center. LGII: *ufd-2(tm1380)*. LGIV: *lin-45(n2018), lin-45(n2506)*. LGV: *sel-10(ok1632)*. The RNAi-sensitized mutant *nre-1(hd20) lin-15B(hd126)* displays enhanced RNAi effectiveness in VPCs (*47*). The transgenes *arIs222 [lag-2p::2xNLS-tagRFP::unc-54 3′UTR]* (*26*) and *arTi4 [lin-31p::YFP-lin-45(417-480)]* (*28*) were previously described.

### *Plasmids and transgenes used for expression in* C. elegans

Plasmids used for transgenes are described in Table S2. *lin-45* transgenes contain *lin-45* cDNA (Wormbase sequence Y73B6A.5a); mutations were generated using PCR and cloned by Gibson assembly. For YFP reporters, the YFP and *lin-45* coding sequences were fused in frame to produce a sequence encoding N-terminally tagged YFP-LIN-45 protein. The *ufd-2* transgene contains *ufd-2* cDNA (Wormbase sequence T05H10.5b.1). For expression in VPCs, coding sequences were cloned into pCC395, a vector derived from pCFJ910 (*48*), containing the visible pharynx marker *myo-2p::tagRFP* and regulatory elements of the *lin-31* gene (*49*) and a 3′ UTR derived from the *unc-54* gene. Alternatively, some transgenes were cloned into pCC249 (de la Cova, et al. 2020), which has identical sequences but lacks *myo-2p::tag-RFP*.

Single-copy miniMos insertion transgenes were generated by germline injection of N2 strain hermaphrodites with 10 ng/μL transgene plasmid and a co-injection mixture of pCFJ601, pCFJ90, pGH8, and pMA122; strains carrying integrated reporters were isolated as described previously (*48*).

### C. elegans *alleles made by gene editing*

Allele *cov37* contains a *GFP-3xFLAG* insertion at the start of *lin-45* isoform Y73B6A.5a, resulting in an N-terminal fusion encoding GFP-3xFLAG-LIN-45. Allele *cov40* contains a *GFP-3xFLAG* insertion at the stop codon shared by all *lin-45* isoforms, resulting in a C-terminal fusion encoding LIN-45-3xFLAG-GFP. All guide RNAs used in this work were designed using the software SapTrap Builder (*50*) and are listed in Table S3. The repair templates plasmids contain *GFP-3xFLAG* and a self-excising cassette and are derived from pDD282 (*51*). pKS4 (for *cov37*) contains two 700bp genomic fragments that flank the *lin-45* start codon; the plasmid pAD2 (for *cov40*) contains 700bp fragments that flank the *lin-45* stop codon.

Gene-edited alleles were generated by germline injection of N2 strain hermaphrodites with 10 ng/μL of repair template plasmid and a ribonucleoprotein mixture as described previously (*52*), specifically: 250 ng/μL Cas9 protein (Macrolabs, University of California Berkeley), 100 ng/μL Alt-R-Crispr-Cas9 tracrRNA (Integrated DNA Technologies catalog 1072532), and 55 ng/μL custom gene-specific Alt-R crRNA synthesized by Integrated DNA Technologies, 2.5 ng/μL pCFJ90, and 20 ng/μL pGH8. Isolation of new gene-edited strains was performed as described previously (*51*), and alleles were validated by PCR and Sanger DNA sequencing of the GFP knock-in regions.

#### Assessment of Multivulva and lethality phenotypes

To assess the Multivulva phenotype, L4 hermaphrodites from uncrowded cultures were picked to fresh plates; adults were examined ∼24 h later using a dissecting microscope and scored for the presence of a normal vulva and the number of any pseudovulvae. All *lin-45(+)* and *lin-45(V627E)* animals scored for the Multivulva phenotype were grown at 20°C.

To assess L1 lethality of *lin-45* hypomorphic mutants, L4 hermaphrodites were picked individually and transferred to fresh plates daily. At ∼48 h after egg laying, all progeny were examined using a dissecting microscope and scored as live larvae, dead larvae, or unhatched eggs. All genotypes for the *lin-45(n2018)* experiment carried the mutation *dpy-20(e1282)*; all genotypes for the *lin-45(n2506)* experiment carried the mutation *unc-24(e138)*.

#### Imaging of VPCs

For imaging of VPCs during larval development, larvae were mounted on an agarose pad in M9 buffer containing 10 mM levamisole. Unless otherwise specified, adults were allowed to lay eggs for 12-16 hours and larvae were grown at 20°C for approximately 2 days after egg collections, a time point when larvae of early, mid and late L3 stage were easily found. To further refine the developmental stage, only L3 stage larvae in which it was evident that VPCs had undergone either one or two cell divisions were scored and reported here.

Qualitative scoring of *lag-2* reporter expression was performed using a Zeiss Axio Imager microscope equipped with widefield LED fluorescent illumination (Excelitas Technologies) and CMOS camera (Zeiss). The locations of VPC descendants after division were determined using DIC optics. Images of RFP expression from *arIs222* [*lag-2p::2xNLS-tagRFP*] were acquired using 500 ms exposure and 100% LED power. Expression was scored on a positive/negative basis. Descendants of P3.p, P4.p, and P8.p which had divided and fused with the hypodermis were scored as negative.

Images used for quantitation were acquired using a Nikon Ti inverted microscope equipped with a spinning disk confocal system (Crest Optics) and dual sCMOS cameras (Teledyne Photometrics). To quantify YFP-tagged LIN-45, Z-stacks of YFP were acquired using a 40x objective; for all images, an exposure time of 700 ms and laser power of 25% was used. To assess ERK-KTR localization in VPCs before cell division in the early L3 stage, animals were synchronized using 2-hour egg collections, grown at 25°C, and imaged at ∼28 hours after egg collection. For all ERK-KTR experiments, Z-stacks of mClover and mCherry were acquired using dual cameras and a 60x objective; for all images, exposure time and laser power used for mClover and mCherry were equal: 500 ms and 25% laser power. Blank images were acquired using the same parameters.

#### RNAi treatment

Plasmids used in RNAi knock-down experiments are derived from the RNAi feeding vector L4440 (Addgene plasmid 1654). The *lacZ* RNAi plasmid was a gift from Iva Greenwald. The *sel-10* RNAi plasmid contains full *sel-10* coding sequence (Wormbase sequence F55B12.3a.1). The *cdc-48*.*1* RNAi plasmid contains a 1073 bp fragment of *cdc-48*.*1* sequence. The *cdc-48*.*2* RNAi plasmid contains a 716 bp fragment of *cdc-48*.*2* sequence. For both *cdc-48* plasmids, fragments of genomic DNA were amplified using primers listed in Table S4 and cloned in the vector L4440 by Gibson assembly.

For each RNAi condition, an HT115(DE3) *E. coli* strain carrying the relevant RNAi plasmid was grown in liquid culture and spread on NGM plates containing IPTG, as previously described (*53*). Simultaneous knock-down of *cdc-48*.*1* and *cdc-48*.*2* was achieved by plating a 1:1 mixture of two strains, carrying *cdc-48*.*1* and *cdc-48*.*2* plasmids. All *C. elegans* genotypes treated with RNAi carried the mutations *nre-1(hd20) lin-15B(hd126), and* were synchronized by a standard bleach/sodium hydroxide protocol to prepare eggs (*54*). Approximately 200 eggs were placed on each plate and grown at 20°C. L3 stage larvae were examined ∼44 hours after egg preparation.

#### Image quantitation and analysis

The subcellular localization of the ERK-KTR biosensor was quantified using methods described in de la Cova, et al. (2017) and the Nikon NIS-Elements software. Briefly, illumination correction and background subtraction was performed using the blank images obtained during each experiment. Image segmentation followed by manual curation was performed in NIS-Elements to create regions of interest (ROIs) for the nucleus (excluding the nucleolus) and cytoplasm of VPCs P4.p, P5.p, P6.p, P7.p, and P8.p. (P3.p fuses with the hyp7 syncytium in approximately half of animals and was not analyzed.) Mean fluorescence intensity for mClover within the nucleus and cytoplasm ROIs was determined for up to five of the most equatorial Z slices per cell. Data presented is a ratio of the mean cytoplasmic mClover intensity/mean nuclear mClover intensity, referred to as “Cyto/Nuc” ratio, per individual cell.

Expression of YFP-tagged LIN-45 protein reporters was quantified using NIS-Elements software. ROIs representing the cytoplasm of P5.pxx, P6.pxx, and P7.pxx cells were created manually using NIS-Elements software. While all other YFP-LIN-45 reporters were exclusively cytoplasmic, the truncated YFP-LIN-45(417-480) form was both cytoplasmic and nuclear. For this reason, the nucleus was included in the quantification of this mutant. Data presented is the mean YFP fluorescence intensity for the most equatorial Z slice, per set of VPC descendants P5.pxx, P6.pxx, and P7.pxx.

#### Statistical analysis

All statistical analyses were performed using GraphPad Prism software. For qualitative scoring of Muv, L1 lethal, *lag-2* expression, and GFP-LIN-45 protein accumulation, we compared the frequencies between two groups using Fisher’s exact test to calculate a two-tailed *P*-value.

For quantitative measurements of ERK-KTR Cyto/Nuc ratio, we compared the means of multiple groups by performing a one-way ANOVA, followed by planned multiple comparisons for the same VPC between different genotypes. (For example, the mean for P6.p of *+; lin-45(+)* larvae was compared to the mean for P6.p of *ufd-2(0); lin-45(+)* larvae.) The reported two-tailed *P*-values were adjusted using Sidak’s multiple comparisons test.

For quantitative measurements of YFP intensities, we compared the means of multiple groups by performing a Welch ANOVA; this test was used because the variance of YFP intensity differed greatly between different genotypes. Our analysis was followed by multiple comparisons between all groups and the reported two-tailed *P*-values were adjusted using Dunnett’s multiple comparisons test.

## Supplementary Materials

**Fig. S1.**
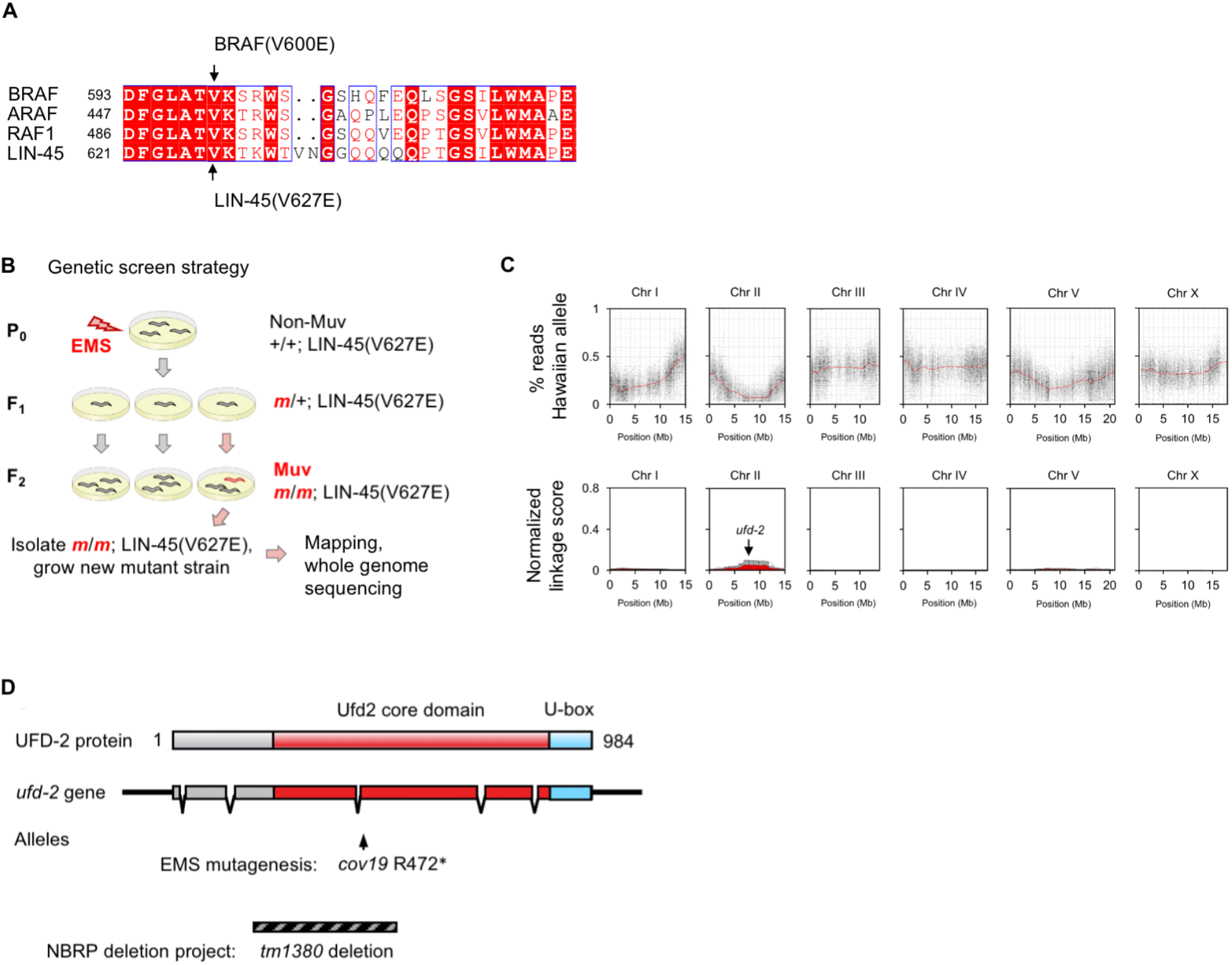
The *lin-45(V627E)* mutation and genetic screen for enhancers. **A)** Alignment of the kinase activation segment in human BRAF, RAF1, ARAF, and *C. elegans* LIN-45. Indicated are the valine mutated in *BRAF(V600E)* and *lin-45(V627E)* (arrows). **B)** Diagram of EMS mutagenesis screen for recessive mutations that enhance the Muv phenotype of *lin-45(V627E)*. **C)** Linkage data from crosses of Muv mutant *cov19; lin-45(V627E)* and Hawaiian wild-type isolate CB4856. Percentage of sequencing reads bearing the Hawaiian variant versus chromosome location (top panel) is shown for pooled F2 progeny that are Muv and presumably homozygous for the recessive *cov19* mutation. Normalized linkage score versus chromosome location (bottom panel). Red and gray columns within Chromosome 2 indicate regions significantly linked to the *cov19* Muv phenotype. The location of the *ufd-2* gene is indicated (arrow). **D)** Diagram of the UFD-2 protein and the *ufd-2* gene. The allele *cov19* is a single nucleotide substitution resulting in a nonsense change at codon 472. The allele *tm1380* obtained from the NRBP deletion project causes a frameshift and premature stop.

**Fig. S2.**
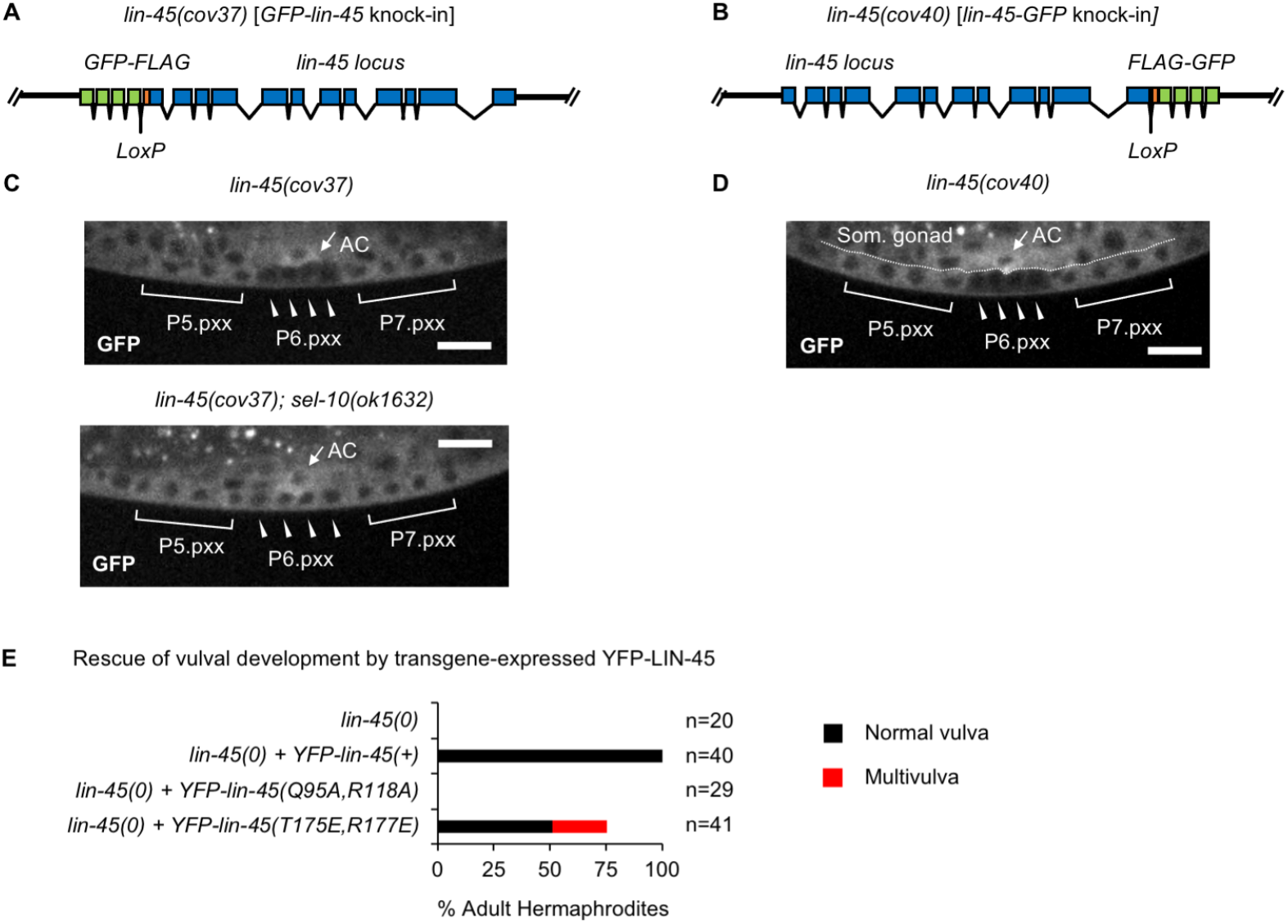
LIN-45 protein in VPCs. **A)** Diagram of the *lin-45(cov37)* knock-in allele, encoding N-terminally tagged GFP-3xFLAG-LIN-45. **B) T**he *lin-45(cov40)* knock-in allele, encoding C-terminally tagged LIN-45-3xFLAG-GFP. **C)** GFP-LIN-45, in *lin-45(cov37)* control (top panel) the *lin-45(cov37); sel-10(0)* mutants (bottom panel) at the L3 larval stage. In all images, descendants of P5.p, P6.p, and P7.p are labeled P5.pxx, P6.pxx, and P7.pxx. Scale bar, 10 μm. **D)** LIN-45-GFP, in *lin-45(cov40)* at the L3 larval stage. **E)** The *lin-45* null mutant *lin-45(dx19)*, referred to as *lin-45(0)*, lacks adult vulva structures. Percentage of *lin-45(0)* adult hermaphrodites that display a normal vulva (black) phenotype is shown for the null mutant alone (n=20), and the null mutant carrying either *YFP-lin-45(+)* (n=40), or *YFP-lin-45(Q95A, R118A*) (n=29), or *YFP-lin-45(T175E, R177E)* (n=41). In addition, adults carrying *YFP-lin-45(T175E, R177E)* displayed the Muv phenotype (red) characteristic of hyperactive Raf-MEK-ERK signaling.

**Fig. S3.**
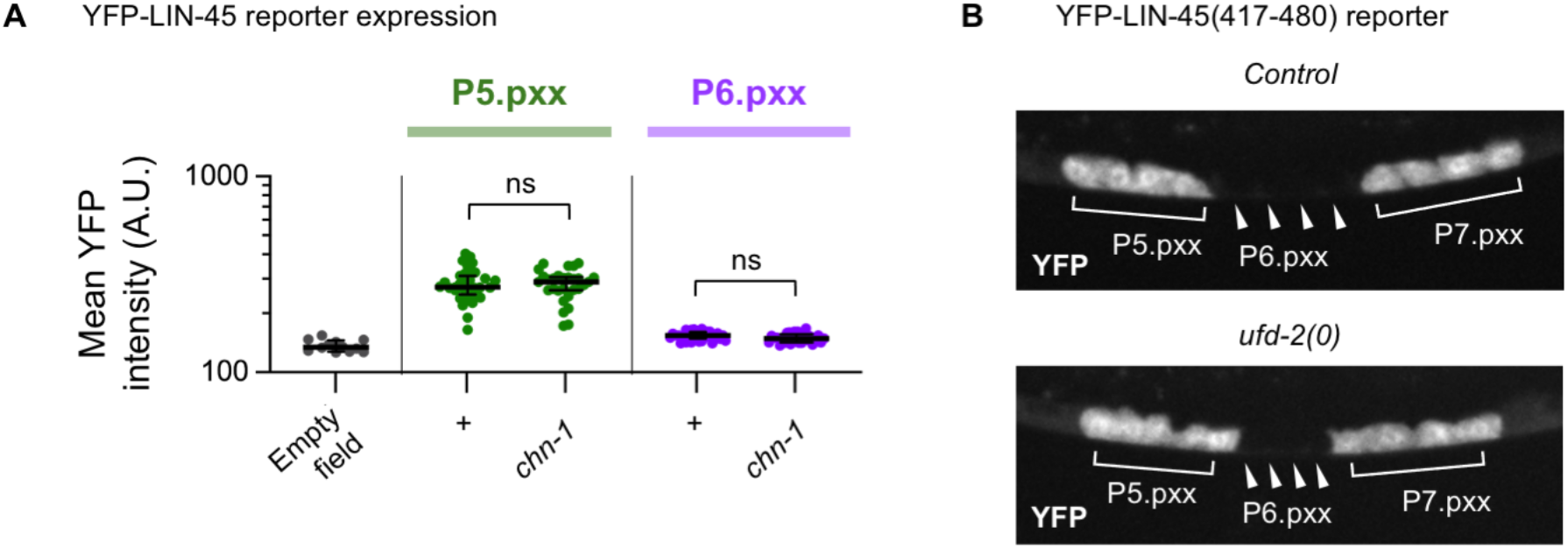
Tests of *chn-1* and *cdc-48* roles in LIN-45 degradation. **A)** Expression of YFP-tagged LIN-45 in P5.pxx cells (green) and P6.pxx cells (purple). To display data from bright and dim cells together, the mean YFP intensities are plotted on a log_10_ scale. YFP-LIN-45 intensity in wild type (n=29) and *chn-1(0)* (n=28) mutants. To test for significant differences in YFP intensity, a Welch ANOVA test was performed. ns indicates not significant. **B)** A short substrate termed the “minimal degron” consists of a YFP-tagged LIN-45 residues 417-480, a region containing the CPD motif. YFP-LIN-45(417-480) reporter expression in L3 stage larvae of wild-type (control) or *ufd-2(0)* mutants. In all images, descendants of P5.p, P6.p, and P7.p are labeled P5.pxx, P6.pxx, and P7.pxx, respectively. ns not signficant.

**Fig. S4.**
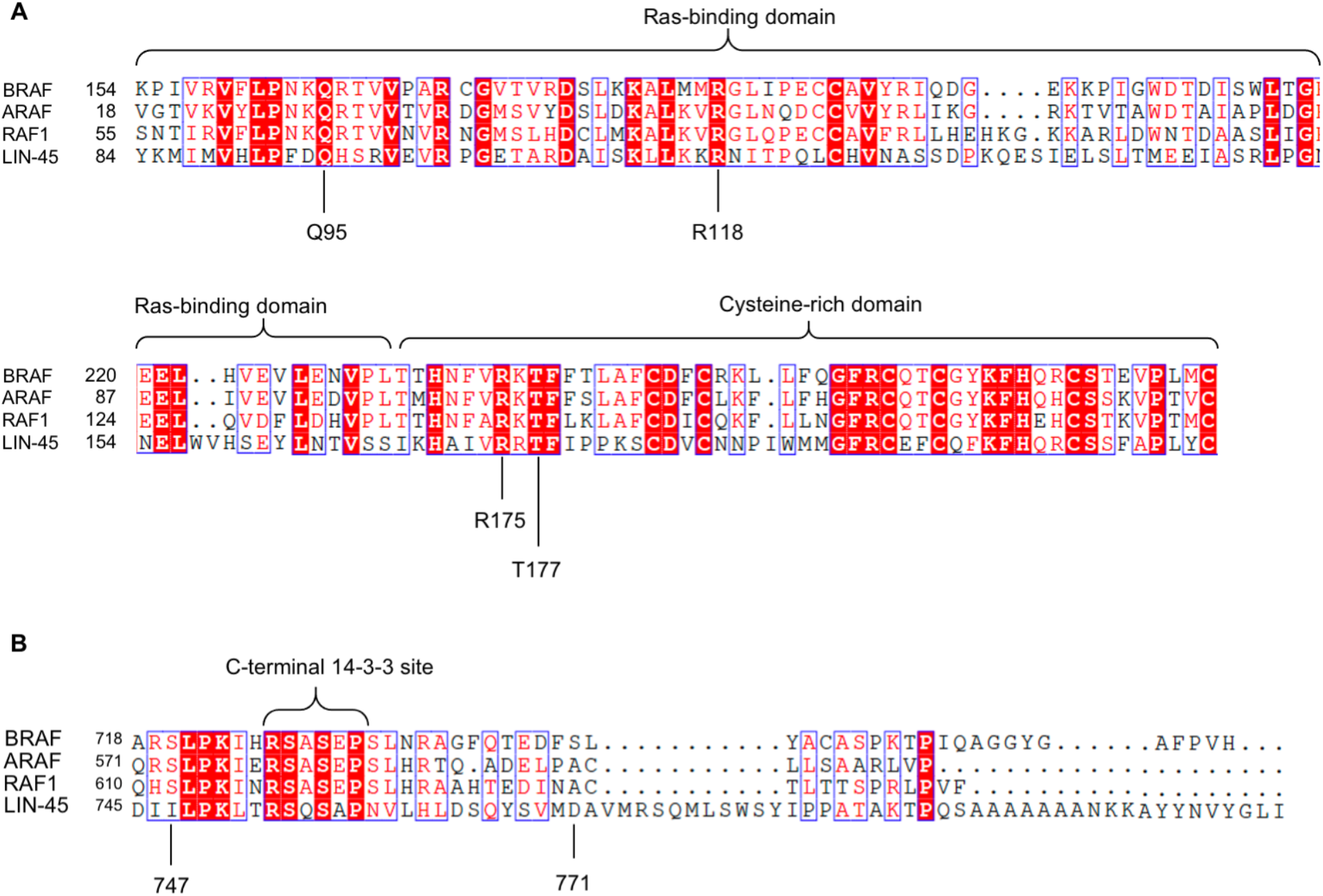
Conserved regions of LIN-45 required for UFD-2 dependence. Alignments show protein sequence of human Raf proteins BRAF, ARAF, RAF1, and *C. elegans* LIN-45. **A)** The N-terminal Ras-binding domain (155-232) and cysteine-rich domain (233-280). The sites of mutations in the RBD (Q95A, R118A), and sites in the CRD (R175, T177) are indicated. **B)** The C-terminal 14-3-3 binding site. The endpoints of two truncations tested (I747 and A772) are indicated.

**Table S1.** *C. elegans* strains used in this work.

| **Strain** | **Genotype** | **Figure** |
| --- | --- | --- |
| N2 | Wild-type isolate | 1 |
| PP198 | *ufd-2(tm1380)* | 1 |
| MKE339 | *covTi106 [lin-31p::lin-45(V627E)::unc-54 3'UTR]* | 1 |
| MKE341 | *ufd-2(tm1380); covTi106 [lin-31p::lin-45(V627E)::unc-54 3'UTR]* | 1 |
| MKE128 | *covTi36 [lin-31p::ufd-2::unc-54 3'UTR]* | 1 |
| MKE372 | *covTi36 [lin-31p::ufd-2::unc-54 3'UTR]; covTi106 [lin-31p::lin-45(V627E)::unc-54 3'UTR]* | 1 |
| MKE343 | *covTi106 [lin-31p::lin-45(V627E)::unc-54 3'UTR]; sel-10(ok1632)* | 1 |
| MKE392 | *ufd-2(tm1380); covTi106 [lin-31p::lin-45(V627E)::unc-54 3'UTR]; sel-10(ok1632)* | 1 |
| MKE236 | *ufd-2(tm1380); dpy-20(e1282)* | 1 |
| WU48 | *lin-45(n2018) dpy-20(e1282)* | 1 |
| MKE237 | *ufd-2(tm1380); lin-45(n2018) dpy-20(e1282)* | 1 |
| MKE242 | *ufd-2(tm1380); unc-24(e138)* | 1 |
| WU49 | *lin-45(n2506) unc-24(e138)* | 1 |
| MKE241 | *ufd-2(tm1380); lin-45(n2506) unc-24(e138)* | 1 |
| GS7090 | *arls222 [lag-2p::2xNLS-tagRFP]* | 2 |
| MKE111 | *ufd-2(tm1380); arls222 [lag-2p::2xNLS-tagRFP]* | 2 |
| MKE363 | *covTi107 [lin-31p::lin-45(V627E)::unc-54 3'UTR]; arls222 [lag-2p::2xNLS-tagRFP]* | 2 |
| MKE365 | *ufd-2(tm1380); covTi107 [lin-31p::lin-45(V627E)::unc-54 3'UTR]; arls222 [lag-2p::2xNLS-tagRFP]* | 2 |
| MKE246 | *covTi15 [lin-31p::ERK-KTR-mClover-T2A-mCherry-his-58::unc-54 3'UTR]* | 2 |
| MKE248 | *ufd-2(tm1380); covTi15 [lin-31p::ERK-KTR-mClover-T2A-mCherry-his-58::unc-54 3'UTR]* | 2 |
| MKE387 | *covTi15 [lin-31p::ERK-KTR-mClover-T2A-mCherry-his-58::unc-54 3'UTR]; covTi107 [lin-31p::lin-45(V627E)::unc-54 3'UTR]* | 2 |
| MKE388 | *ufd-2(tm1380); covTi15 [lin-31p::ERK-KTR-mClover-T2A-mCherry-his-58::unc-54 3'UTR]; covTi107 [lin-31p::lin-45(V627E)::unc-54 3'UTR]* | 2 |
| MKE277 | *lin-45(cov40)* | 3, S2 |
| MKE243 | *lin-45(cov37)* | 3 |
| MKE267 | *ufd-2(tm1380); lin-45(cov37)* | 3 |
| MKE291 | *lin-45(cov37); sel-10(ok1632)* | 3, S2 |
| MKE232 | *covTi64 [lin-31p::yfp-lin-45::unc-54 3'UTR]* | 3 |
| MKE234 | *ufd-2(tm1380); covTi64 [lin-31p::yfp-lin-45::unc-54 3'UTR]* | 3 |
| MKE266 | *covTi64 [lin-31p::yfp-lin-45::unc-54 3'UTR]; sel-10(ok1632)* | 3 |
| MKE440 | *ufd-2(tm1380); covTi64 [lin-31p::yfp-lin-45::unc-54 3'UTR]; sel-10(ok1632)* | 3 |
| MKE496 | *covTi169 [lin-31p::yfp-lin-45(T432A,S436A)::unc-54 3'UTR]* | 3 |
| MKE497 | *ufd-2(tm1380); covTi169 [lin-31p::yfp-lin-45(T432A,S436A)::unc-54 3'UTR]* | 3 |
| MKE441 | *covTi126 [lin31p::yfp-lin-45(V627E)]* | 3 |
| MKE442 | *ufd-2(tm1380); covTi126 [lin31p::yfp-lin-45(V627E)]* | 3 |
| GS7738 | *arTi4 [lin-31p::yfp-lin-45(417-480)::unc-54 3'UTR]* | 4 |
| MKE142 | *ufd-2(tm1380); arTi4 [lin-31p::yfp-lin-45(417-480)::unc-54 3'UTR]* | 4 |
| MKE422 | *covTi64[lin-31p::yfp-lin-45::unc-54 3'UTR]; nre-1(hd20) lin-15B(hd126)* | 4 |
| GS7825 | *arTi4 [lin-31p::yfp-lin-45(417-480)::unc-54 3'UTR]; nre-1(hd20) lin-15B(hd126)* | 4 |
| MKE382 | *covTi110 [lin-31p::yfp-lin-45(288-813)::unc-54 3'UTR]* | 5 |
| MKE384 | *ufd-2(tm1380); covTi110 [lin-31p::yfp-lin-45(288-813)::unc-54 3'UTR]* | 5 |
| MKE463 | *covTi159[lin-31p::yfp-lin-45(167-813)::unc-54 3'UTR]* | 5 |
| MKE464 | *ufd-2(tm1380); covTi159[lin-31p::yfp-lin-45(167-813)::unc-54 3'UTR]* | 5 |
| MKE484 | *covTi165 [lin-31p::yfp-lin-45(82-813)::unc-54 3'UTR]* | 5 |
| MKE485 | *ufd-2(tm1380); covTi165 [lin-31p::yfp-lin-45(82-813)::unc-54 3'UTR]* | 5 |
| MKE181 | *covTi54 [lin-31p::yfp-lin-45(1-747)::unc-54 3'UTR]* | 5 |
| MKE192 | *ufd-2(tm1380); covTi54 [lin-31p::yfp-lin-45(1-747)::unc-54 3'UTR]* | 5 |
| MKE426 | *covTi127[lin-31p::yfp-lin-45(1-772)::unc-54 3'UTR]* | 5 |
| MKE427 | *ufd-2(tm1380); covTi127[lin-31p::yfp-lin-45(1-772)::unc-54 3'UTR]* | 5 |
| MKE436 | *covTi158 [lin-31p::yfp-lin-45(Q95A, R118A)::unc-54 3'UTR]* | 5 |
| MKE437 | *ufd-2(tm1380); covTi158 [lin-31p::yfp-lin-45(Q95A, R118A)::unc-54 3'UTR]* | 5 |
| MKE469 | *covTi163[lin-31p::yfp-lin-45(R175E, T177E)::unc-54 3'UTR]* | 5 |
| MKE470 | *ufd-2(tm1380); covTi163[lin-31p::yfp-lin-45(R175E, T177E)::unc-54 3'UTR]* | 5 |
| MKE471 | *covTi62[lin-31p::yfp-lin-45::unc-54 3'UTR]; lin-45(dx19)/nT1; +/nT1[umnlS28]* | S2 |
| MKE473 | *covTi154[lin-31p::yfp-lin-45(Q95A, R118A)::unc-54 3'UTR]; lin-45(dx19)/nT1; +/nT1[umnlS28]* | S2 |
| MKE498 | *covTi163[lin-31p::yfp-lin-45(R175E, T177E)::unc-54 3'UTR]; lin-45(dx19)/nT1; +/nT1[umnlS28]* | S2 |
| EJ521 | *lin-45(dx19)/nT1 [unc(n754) let] (IV;V)* | S2 |
| MKE233 | *chn-1(by155); covTi64 [lin-31p::yfp-lin-45::unc-54 3'UTR]* | S3 |

**Table S2.** Plasmids used in this work.

| **Plasmid** | **Description** | **Source** |
| --- | --- | --- |
| pCCRT23 | *miniMos lin-31p::lin-45(V627E)::unc-54 3'UTR, myo-2p::tagRFP::unc-54 3'UTR* | This work |
| pAD1 | *miniMos lin-31p::ufd-2::unc-54 3'UTR* | This work |
| pCC324 | *miniMos lin-31p::ERK-KTR-mClover-T2A-mCherry-his-58::unc-54 3'UTR* | de la Cova, et al. 2017 |
| pCCRT15 | *miniMos lin-31p::yfp-lin-45::unc-54 3'UTR, myo-2p::tagRFP::unc-54 3'UTR* | This work |
| pCCRT69 | *miniMos lin-31p::yfp-lin-45(T432A, S436A)::unc-54 3'UTR, myo-2p::tagRFP::unc-54 3'UTR* | This work |
| pCCRT45 | *miniMos lin31p::yfp-lin-45(V627E), myo-2p::tagRFP::unc-54 3'UTR* | This work |
| pCC194 | *miniMos lin-31p::yfp-lin-45(417-480)::unc-54 3'UTR* | de la Cova, et al. 2020 |
| pCCRT34 | *miniMos lin-31p::yfp-lin-45(288-813)::unc-54 3'UTR, myo-2p::tagRFP::unc-54 3'UTR* | This work |
| pCCRT55 | *miniMos lin-31p::yfp-lin-45(167-813)::unc-54 3'UTR* | This work |
| pCCRT62 | *miniMos lin-31p::yfp-lin-45(82-813)::unc-54 3'UTR* | This work |
| pCCRT6 | *miniMos lin-31p::yfp-lin-45(1-747)::unc-54 3'UTR, myo-2p::tagRFP::unc-54 3'UTR* | This work |
| pCCRT35 | *miniMos lin-31p::yfp-lin-45(1-772)::unc-54 3'UTR, myo-2p::tagRFP::unc-54 3'UTR* | This work |
| pCCRT46 | *miniMos lin-31p::yfp-lin-45(Q95A, R118A)::unc-54 3'UTR, myo-2p::tagRFP::unc-54 3'UTR* | This work |
| pCCRT56 | *miniMos lin-31p::yfp-lin-45(R175E, T177E)::unc-54 3'UTR* | This work |
| pKS4 | *Repair template for N-terminal GFP-lin-45 knock-in, self-excising cassette* | This work |
| pAD2 | *Repair template for C-terminal lin-45-GFP knock-in, self-excising cassette* | This work |

**Table S3.**
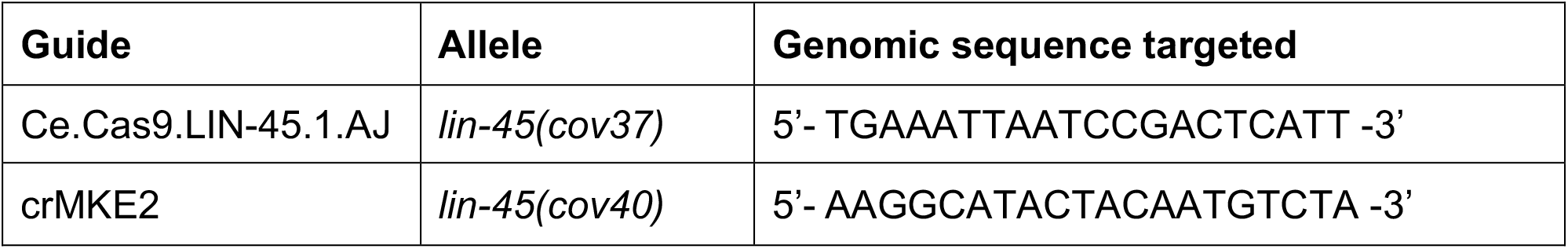
Gene-specific guide RNAs used in this work.

**Table S4.**
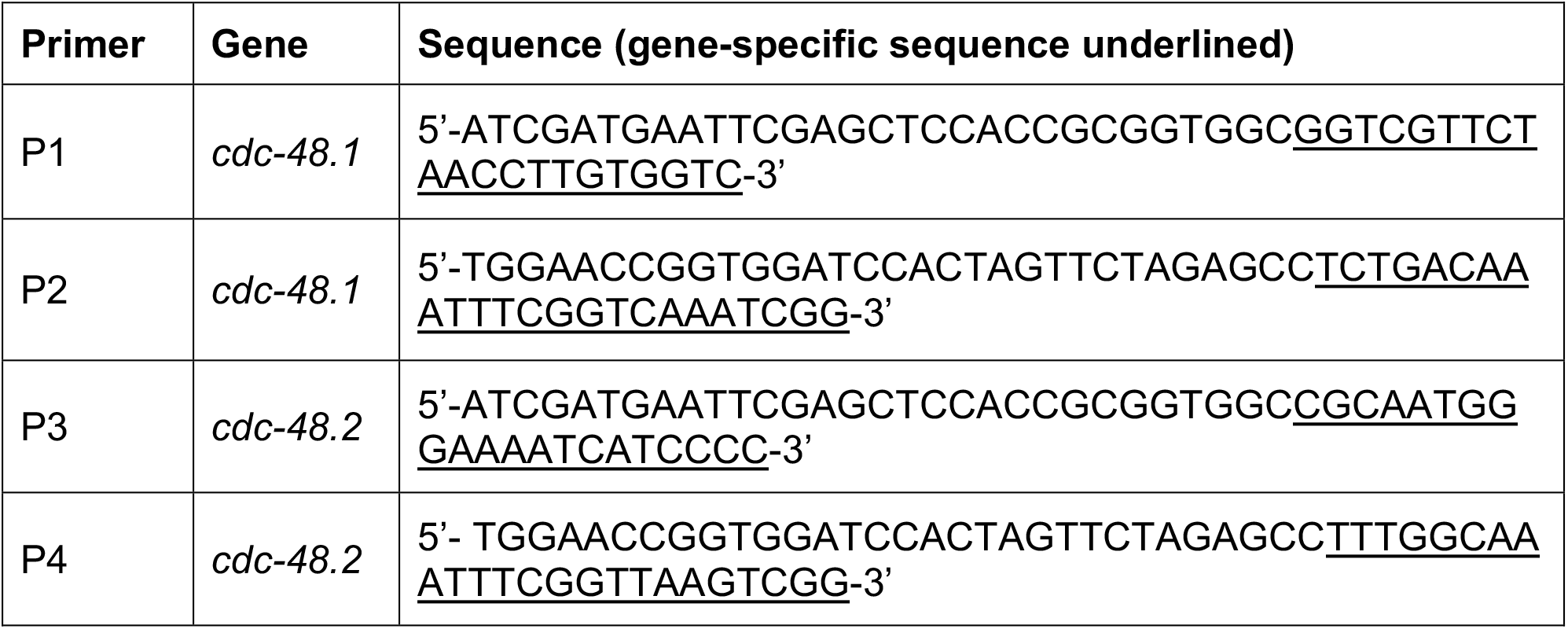
Primers for RNAi used in this work.

## Acknowledgments

The authors thank Hannes Bülow, Jennifer Gutzman, and Christopher Quinn for their comments and discussions.

## Author contributions

Conceptualization: RT, CCD

Formal analysis: RT, AD, CSRT, CCD

Investigation: RT, AD, KSS, CSRT, FC, CCD

Writing: RT, AD, CCD

## Data and materials availability

All data are available in the main text or the supplementary materials.

